# Genetic indicators of drug resistance in the highly repetitive genomes of *Trichomonas vaginalis* and other trichomonads

**DOI:** 10.1101/076729

**Authors:** Martina Bradic, Sally D. Warring, Grace E. Tooley, Paul Scheid, William E. Secor, Kirkwood M. Land, Po-Jung Huang, Ting-Wen Chen, Chi-Ching Lee, Petrus Tang, Steven A. Sullivan, Jane M. Carlton

## Abstract

**Background:** *Trichomonas vaginalis*, the most common non-viral sexually transmitted parasite, causes ~283 million trichomoniasis infections annually and is associated with complications during pregnancy and increased risk of HIV-1 acquisition. The antimicrobial drug metronidazole is used for treatment, but has lead to widespread drug resistance. We undertook sequencing of multiple clinical isolates and lab derived lines to identify genetic markers and mechanisms of metronidazole resistance.

**Results:** Reduced representation genome sequencings of more than 100 *T. vaginalis* clinical isolates identified 3,923 SNP markers and presence of a bipartite population structure. Linkage disequilibrium decays rapidly, suggesting genome-wide recombination and the feasibility of genetic association studies in the parasite. We identified 72 SNPs associated with metronidazole resistance, and a comparison of SNPs within several lab-derived resistant lines revealed an overlap with the clinically resistant isolates. We identified SNPs in sets of genes for which no function has yet been assigned, as well as in functionally-characterized genes highly relevant to drug resistance (*e.g., pyruvate:ferredoxin oxidoreductase* (*PFO*)). Transcription profiling of these and other genes served as proxy for testing the functional consequences of multiple identified SNPs. Transcription profiles of lab-derived drug resistance strain as well as clinically resistant strain depict common regulation changes in carbohydrate metabolism and oxygen detoxification pathways correlated with Mz resistance. Finally, we identified convergent genetic changes in lab-derived resistant lines of *Tritrichomonas foetus*, a distantly-related species that causes venereal disease in cattle.

**Conclusions:** Our observation of shared genetic changes within and between *T. vaginalis* and *Tr. foetus* parasites suggests conservation of the pathways through which adaptation has occurred. These findings extend our knowledge of drug resistance in the parasite, providing a panel of markers that can be used as a diagnostic tool.

## Background

Trichomonads are haploid unicellular, microaerophilic members of eukaryotic phylum Parabasalia that infect a variety of vertebrates including humans, wildlife, farm animals, and pets [1]. Among the several important species in this group is *Trichomonas vaginalis,* which causes trichomoniasis, the most common non-viral sexually transmitted disease in humans, with 3.7 million cases reported each year in the United States [2]. *T. vaginalis* can cause long-term symptomatic infections of vulvar and urethral regions of the genital tract [3]. Importantly, *T. vaginalis* infection is associated with increased risk of HIV infection [4], development of prostate cancer [5], and complications during pregnancy, such as premature and low-weight births [6]. *Tritrichomonas foetus* is an economically important trichomonad cattle parasite which, though only distantly related to *T. vaginalis*, nevertheless has an analogous site of infection and disease etiology [7–9].

A highly fragmented assembly of the ~160 Mb *T. vaginalis* strain G3 genome generated by Sanger sequencing was published in 2007. It consists of ~17,000 scaffolds containing ~60,000 predicted protein coding genes, many of them members of high copy number gene families, and ~40,000 transposable element (TE) genes distributed among ~59 TE families, members of which are highly similar (average ~97.5%) within a family [10]. Identification of meiosis-specific genes in the genome suggested that the parasite might have a cryptic sexual cycle or may have recently lost the ability to undergo genetic exchange [10, 11]. Genetic markers mined from the genome have been used in population studies of the species [12, 13], revealing significant *T. vaginalis* genetic diversity that is structured but unrelated to geographical origin [14]. To date, a genome assembly for *Tr. foetus* has not been published, though DNA reannealing assays indicate that its genome too is highly repetitive [15].

Metronidazole (Mz) and tinidazole are the only two United States Food and Drug Administration approved drugs effective against *T. vaginalis*. However, clinical failure of Mz treatment has been reported since 1959 [16] and ranges from ~4% in the U.S.A. [17] to 17% in Papua New Guinea [18]. Mz is a pro-drug, which must be reduced at its nitro group in order to form the nitroradical anions that are toxic to the parasite. The electrons required for Mz activation are generated through several mechanisms, including the reduction of pyruvate by the activity of *pyruvate:ferredoxin oxidoreductase* (*PFO*) [19-22], an enzyme found in membrane-bound organelles called hydrogenosomes that produce ATP and hydrogen, as well as several pathways hypothesized to exist in the cytosol [23-25]. *Tr. foetus* is here again similar to *T. vaginalis* in displaying both Mz resistance and related hydrogenosomal metabolic pathways [24, 26-28].

Aerobic and anaerobic pathways of Mz resistance have been proposed. Clinical (aerobic) resistance is typically observed only in the presence of oxygen, since oxygen decreases Mz toxicity by re-oxidizing nitroradical anions before they can damage the parasite, reverting Mz to an inactive form. Clinically resistant *T. vaginalis* strains have impaired oxygen scavenging activity and elevated levels of intracellular oxygen. Oxygen stress response genes seem to be involved in aerobic Mz resistance, with *NADH-dependent oxidase* and *flavin reductase* (*FR*) in particular being implicated as important enzymes involved in oxygen removal [23, 26]. In contrast, anaerobic resistance has typically been observed only *in vitro* [27, 28], with the exception of strain B7268 where anaerobic resistance developed in a patient [29]. Anaerobically resistant strains are marked by the absence of certain hydrogenosome metabolic enzymes that would normally activate Mz, including *PFO* and *ferredoxin* (*Fd*) [21, 30, 31], as well as cytosolic *thioredoxin reductase* (*TrxR*) [24]. Aerobic and anaerobic resistance pathways are proposed to be fundamentally different, although *FR*, whose primary role is to reduce molecular oxygen to hydrogen peroxide, appears to be involved in both processes [23]. To date, studies of the mechanism of Mz resistance have focused wholly on candidate genes known to be involved in oxygen scavenging or hydrogenosome metabolism, for example, *PFO, Fd, FR,* and *TrxR* [20, 23, 25, 28, 32]. However, alterations in these genes do not explain all cases of Mz resistance, nor have other genes been identified as possible candidates. The aim of our study was to address these issues by large-scale genetic comparison of Mz sensitive and resistant strains.

High-throughput methods for large-scale genetic comparison and analysis of mutational phenotypes are particularly difficult to implement in trichomonads. The highly repetitive nature of the *T. vaginalis* and *Tr. foetus* genomes makes them refractory to conventional short-read genome sequencing. We therefore employed double digest restriction-site associated DNA sequencing (ddRAD-Seq) to genotype 102 *T. vaginalis* isolates from eight countries, the largest sampling of *T. vaginalis* genomes reported to date. We identified single nucleotide polymorphisms (SNPs) in the sequences in order to explore population structure and genome-wide linkage disequilibrium (LD) in the isolates, and provide a basis for a genome-wide association study of Mz genetic indicators. In addition, we compared laboratory-derived Mz resistant isolates with their isogenic parents to identify SNPs that could indicate Mz resistance pathways common to resistant lines. Similarly, because there is no efficient high-throughput transfection system for large-scale assay of mutational phenotypes in *T. vaginalis*, we conducted whole transcriptome analysis of gene regulation in resistant and sensitive isolates, as a proxy for direct functional testing. Finally, to explore the universality of the genetic bases of Mz resistance among trichomonads, we analyzed genetic changes between a parental strain and *in vitro-*derived Mz resistant lines of *Tr. foetus*.

## Results

**Figure S1** in **Additional file 1** depicts the various components of this study. We used ddRAD-Seq to partially sequence genomes of 102 *T. vaginalis* clinical isolates from eight countries. Sequence reads were filtered, aligned, and single nucleotide polymorphisms (SNPs) were identified (**Additional file 2**; see **Materials and Methods** for filtering criteria). Our final set of 3,923 high quality SNPs comprised 1,463 nonsynonymous mutations, 879 silent (synonymous) mutations, 10 nonsense mutations, and 1,571 intergenic mutations.

### Population structure and fast LD decay suggest recombination in *T. vaginalis* genomes

Population structure and linkage disequilibrium (LD) properties play a key role in determining success of mapping genotypes to phenotypes [33]. The existence of genetic subpopulations and the occurrence of LD decay over short distances are strong indicators of genetic recombination and sexual reproduction in a population [34, 35], with implications for correlating genotypes to drug resistance. To characterize population structure, we performed principal components analysis (PCA) of SNPs from 102 *T. vaginalis* isolates (**Figure 1A**). The first two principal components accounted for the highest variation within the sampled isolates and explained 10.1% and 6.1% of the variation, respectively. Similar to published findings, these two PCA axes split the global sample into two subpopulations [14, 36], a structure congruent with sexual reproduction. Population structure did not correspond to the geographical origin of the isolates.

**Figure 1.**
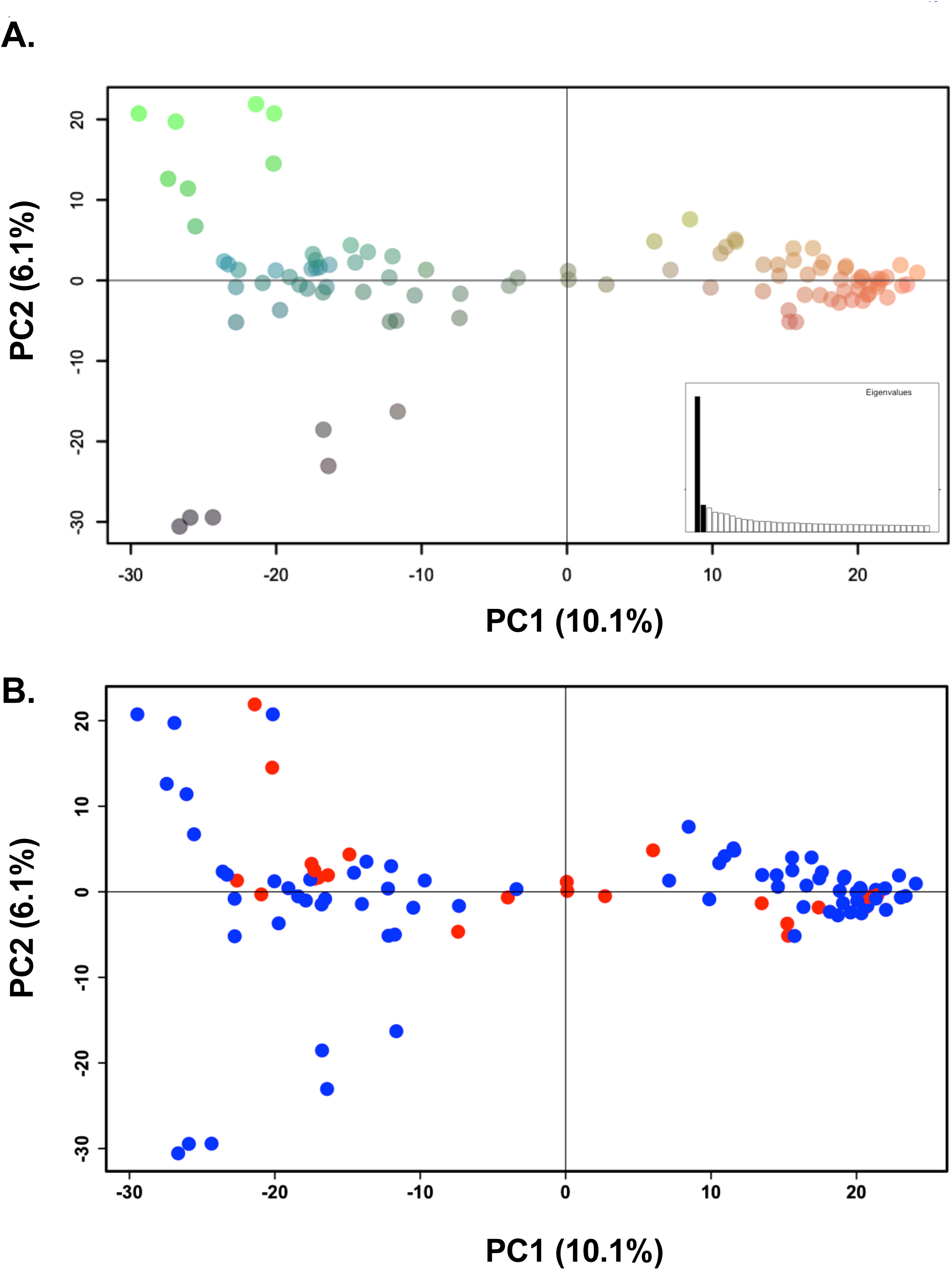
Population structure and Linkage Disequilibrium (LD) in *T. vaginalis* global samples. A. Principal components analysis of 102 *T. vaginalis* isolates using 3,923 genome-wide ddRAD SNPs. The x-axis represents principal component one (PC1) and explains 10.1% of the variation, while the y-axis represents principle component two (PC2), and explains 6.1% of the variation; the inset represents distribution of eigenvalues where each eigenvalue is the variance of the corresponding PC. Each isolate is colored according to a gradient of similarity within each genetic type. **B.** Principal components analysis showing variation in 32 Mz resistant (red) and 63 Mz sensitive (blue) isolates as defined by MLC > 100 ug/mL.

In asexual populations, LD is maintained over long genomic distances or even over the entire genome, while genetic exchange in sexually reproducing populations constrains LD to shorter distances. We plotted LD decay along the 898 genomic contigs containing 2,837 SNPs in one population, and along the 872 contigs containing 2,694 SNPs in the other population, and found decay occurring within 1 kb in both populations (**Figure 2**), although the rate differed between the two. These results, together with the presence of population structure, provide strong evidence that genetic exchange occurs, or occurred sometime in the recent evolutionary past, of both *T. vaginalis* populations. The difference in LD decay rates indicates potential differences in recombination, population size, or phenotypic traits (*e.g*., transmission, drug resistance, virulence) between the two populations.

**Figure 2.**
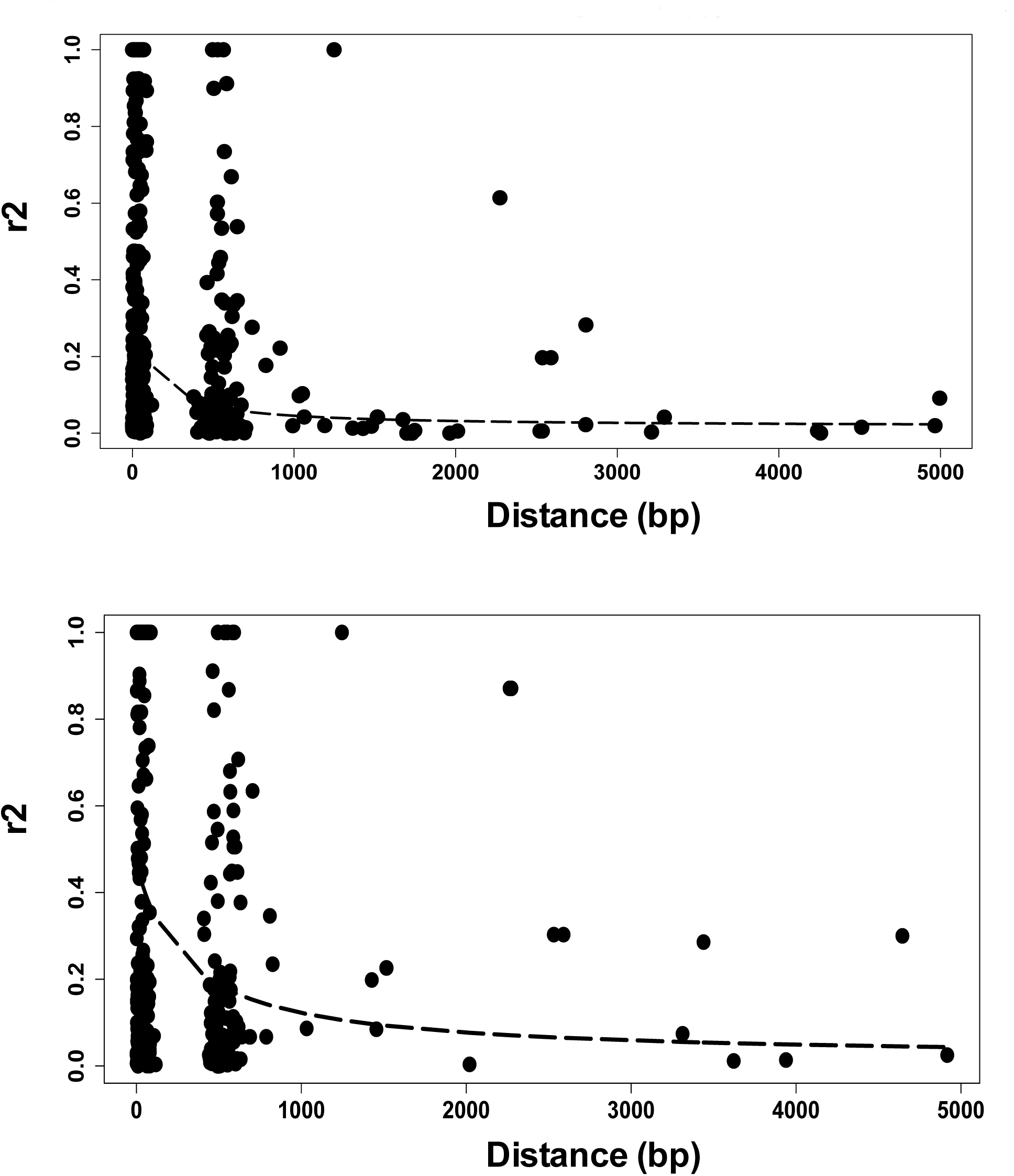
LD decays over distance in two populations of *T. vaginalis*. LD decay calculated over 5 kb intervals in 898 contigs (top panel) and 872 contigs (bottom panel) is shown for the two populations of *T. vaginalis*.

### Association study of *T. vaginalis* isolates identifies candidate biomarkers of Mz resistance

Our findings suggested 1) that population structure should be accounted for in *T. vaginalis* association studies in order to avoid spurious genotype-to-phenotype association; and that 2) fast LD decay allows for mapping of phenotypic traits in candidate regions. We first tested for association between *T. vaginalis* population structure and resistance to metronidazole (Mz) as measured using a standard minimum lethal concentration (MLC) protocol under aerobic conditions (**Additional file 2**). Of the 95 isolates for which MLC could be determined, 63 were classified as 'sensitive' and 32 as 'resistant' (MLC > 100 μg/ml), of which 23 were also ‘highly resistant’ (MLC > 400 μg/ml). Neither of the two subpopulations detected by PCA of ddRAD SNPs was significantly associated with Mz resistance (Pearson's chi-squared test for MLC > 100 μg/ml: χ^2^ = 0.2139, df = NA, p-value = 0.8136; for MLC > 400 μg/ml: χ^2^ = 0.2291, df = NA, p-value = 0.7271; **Figure 1B**).

We next used Discriminant Analysis of Principal Components (DAPC) [37] to identify SNPs that distinguish Mz resistant from Mz sensitive isolates, and quantified the contribution of each allele to this distinction (**Figure 3**). Our analysis showed partial overlap of alleles between sensitive and resistant groups (**Figure 3A**). We identified the SNPs that contribute the most to distinguishing sensitive from resistant phenotypes (**Figure 3B**). DAPC identified a set of 72 SNPs associated with moderate or high Mz resistance (MLC > 100 μg/ml or >400 μg/ml; **Additional file 3**).

**Figure 3.**
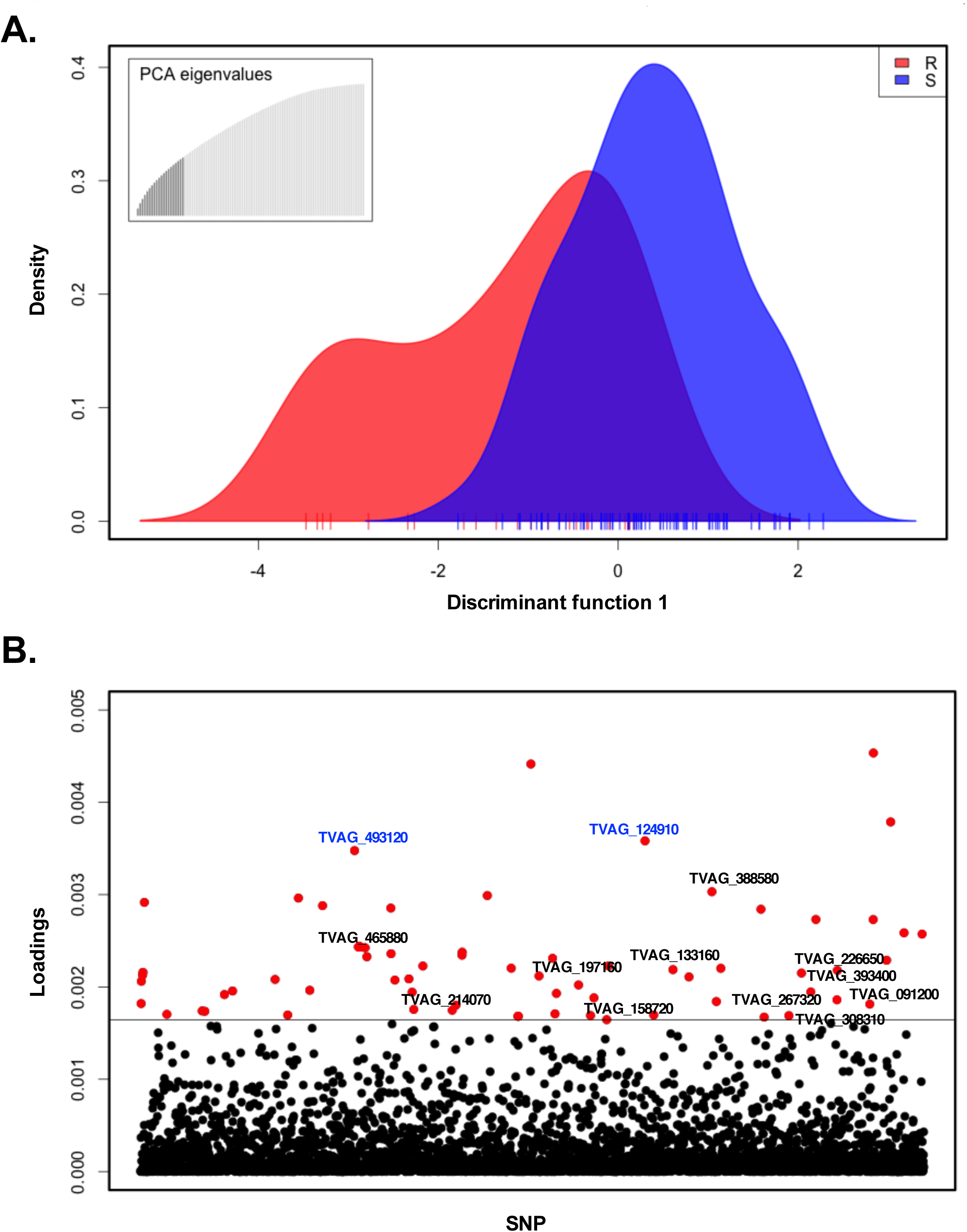
Association mapping for Mz resistance in *T. vaginalis*. **A.** Discriminant analysis of principal components (DAPC) representing variation and distribution of markers between drug resistant (red) and drug sensitive (blue) isolates. Ticks on the x-axis represent individual isolates. The inset represents the distribution of eigenvalues, with the black histogram showing all the principle components that were used in the DAPC analysis. **B.** Loadings plot of the SNPs used for association analysis. The distribution of variances for SNP markers within and between resistant and sensitive isolates is represented as a loading value for each marker. The loading values above the threshold represent the largest between-group variance and the smallest within-group variance and are associated with the resistant phenotype. SNP markers are indicated on the x-axis, and the loadings value for each SNP on the y-axis; a horizontal line indicates the loadings threshold. Red dots represent 72 SNPs identified as significantly contributing to Mz resistance. Two blue-colored dots represent two nonsynonymous SNPs in two conserved hypothetical genes (TVAG_493120 and TVAG_124910) common to both moderate (MLC ≥ 100) and high (MLC ≥ 400 Mz) phenotypes.

These comprised multiple silent (synonymous) and intergenic SNPs, but in addition a total of 16 nonsynonymous SNPs associated with moderate resistance in 13 genes (for example TVAG_197160 CAMK family protein kinase, TVAG_158720 BspA surface antigen), and two nonsynonymous SNPs in two conserved hypothetical genes (TVAG_124910, TVAG_493120) associated with both moderate (MLC ≥ 100 μg/ml) and high (MLC ≥ 400 μg/ml) resistance. A summary of these genes and their SNPs is shown in **Additional file 4.**

### SNP changes in Mz resistant *T. vaginalis* lines derived *in vitro* suggest similar mechanisms of drug resistance

While the 72 SNPs we found associated with moderate or high resistance in disparate clinical isolates represent a potentially important set of genetic markers of the resistance phenotype, such association does not demonstrate causation. To identify SNPs that may be more strongly inferred to be consequences of adaptation to drug treatment, we exploited existing pairs of *T. vaginalis* lines, each consisting of a highly drug-resistant line derived *in vitro* from a drug-sensitive or moderately resistant parental line by increasing drug selection pressure over time (**Additional file 1, Figure S2**). We analyzed SNPs observed in ddRAD-Seq data from three such pairs of parental and derived lines: (1) Mz-sensitive B7708 and highly resistant B7708-M; (2) Mz-sensitive F1623 and highly resistant F1623-M [19]; and (3) moderately resistant B7268 and highly resistant B7268-M [38]. SNPs were determined by comparing the parental and derived line to the reference genome of the Mz-sensitive lab strain G3. For each pair, only bases that were the same in the parental line and the G3 reference but different in the derived strain were considered as SNPs (see **Materials and Methods**). We found 39 SNPs in 36 genes to be common to all three derived, highly resistant lines (**Figure 4**, **Additional file 5)**.

**Figure 4.**
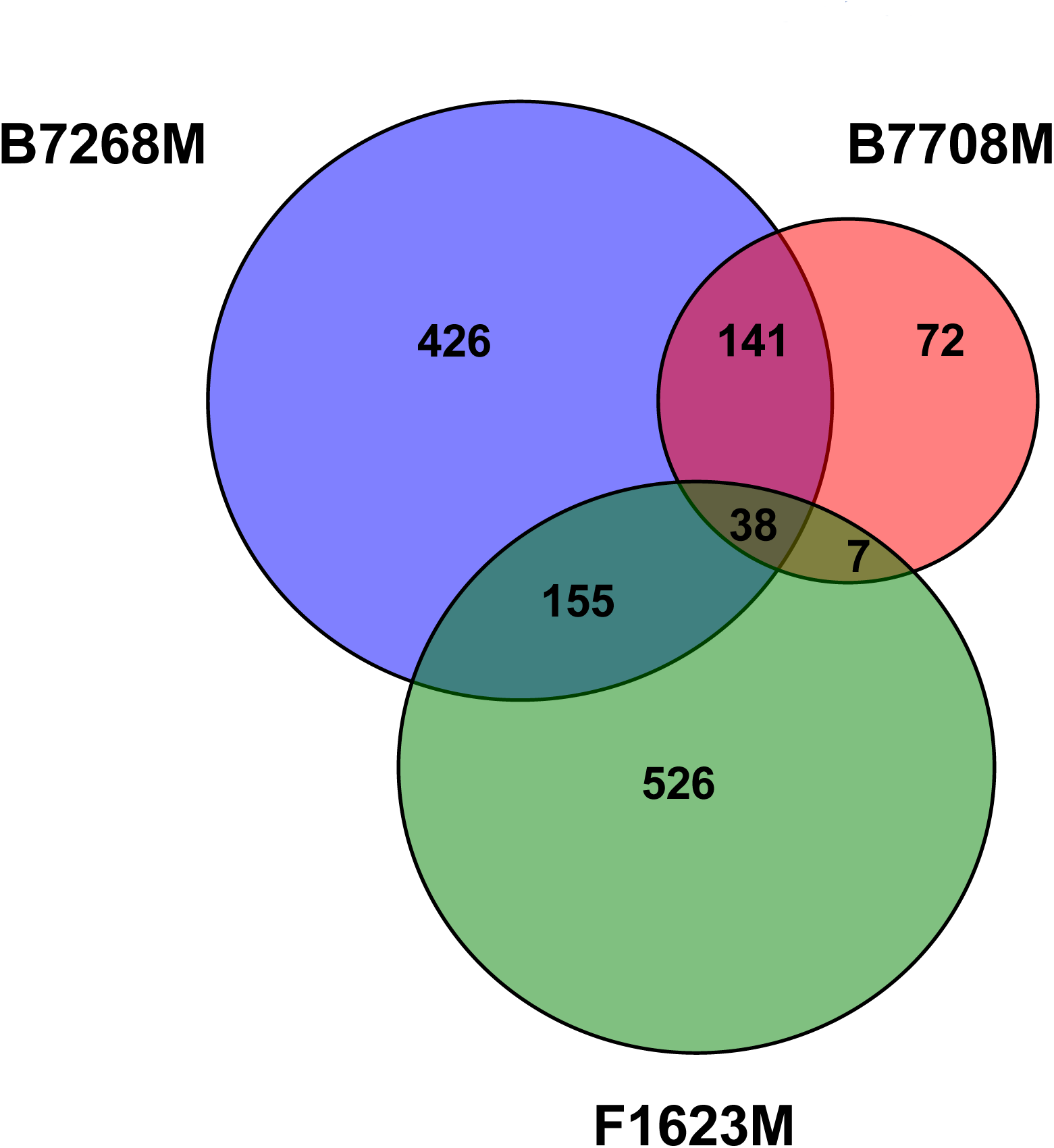
Common genetic changes in laboratory-derived *T. vaginalis* lines. Venn diagram circle sizes are based on numbers of genes that have nonsynonymous SNPs. A nonsynonymous SNP was defined as a position where the nucleotide in the ancestral strain was the same as in the reference G3 strain, but different in the derived (more resistant) strain.

Changes common to two out of three derived lines included seven nonsynonymous mutations in seven genes in B7708-M and F1623-M; 137 nonsynonymous (including one nonsense) SNP in 141 genes in B7708-M and B7268-M; and 170 nonsynonymous and two nonsense SNPs in 155 genes in F1623-M and B7268-M. We also identified multiple synonymous and intergenic SNPs in common between the pairs (data not shown).

Of particular note are five genes (TVAG_124910, TVAG_197160, TVAG_226650, TVAG_393400, TVAG_465880) containing nonsynonymous SNPs that were 1) present in at least two lab-derived resistant strains, and 2) associated with Mz resistance in our association study of ~100 clinical isolates described above. TVAG_197160 encodes a predicted CAMK kinase family protein mentioned above, while the rest encode conserved hypothetical proteins, one of which (TVAG_465880) contains multiple SNPs shared between the derived and clinical isolates. These five genes represent novel candidates for roles in both clinical and *in vitro*-derived Mz resistance. A summary of the genes and their SNPs described in this section is shown in **Additional file 4.**

### Gene expression changes are associated with resistance and SNPs in clinical and derived isolates

We employed whole transcriptome analysis to survey changes in expression of genes associated with resistance as a proxy for direct functional testing of SNP effects. We sequenced RNA-Seq libraries in triplicate from nine Mz-sensitive isolates (the reference strain G3 and GOR69, NYCA04, NYCB20, NYCD15, NYCE32, NYCF20, NYCG31, SD2), and three resistant isolates: the moderately resistant clinical isolate B7268, its highly resistant *in vitro* derivative B7268-M, and highly resistant clinical isolate NYCC37. Of 57,796 predicted genes in the *T. vaginalis* G3 genome (excluding genes predicted as TEs or other highly repeated genes) 28,919 (50%) showed evidence of expression (**Additional file 6)**. Mz resistant strains B7268 and B7268M shared the highest number of pairwise down- and up-regulated genes (**Additional file 6)**, as expected given their common ancestry, and all three resistant strains shared 1,097 up-regulated and 1,212 down-regulated genes (**Figure 5** and **Additional file 6)**. Comparing these genes to results from our two SNP studies described above, we identified three genes that were up- or dowregulated in at least two Mz-resistant lines and also were among the five novel candidate genes flagged in both of the studies: CAMK family protein kinase TVAG_197160 (downregulated in all three resistant lines); conserved hypothetical protein TVAG_226650 (upregulated in two lines, NYCC37 and B7268M); and conserved hypothetical protein TVAG_465880 (upregulated in three lines). We also found three genes that were 1) transcriptionally regulated in at least two resistant strains, and 2) had nonsynonymous SNPs associated with our three lab-derived resistant isolates: TVAG_316160, a Ser/Thr protein phosphatase, which is down-regulated, and conserved hypothetical protein genes TVAG_210010 containing two SNPs and TVAG_019490, both upregulated in two lines. A summary of the genes and their SNPs described in this section is shown in **Additional file 4**.

**Figure 5.**
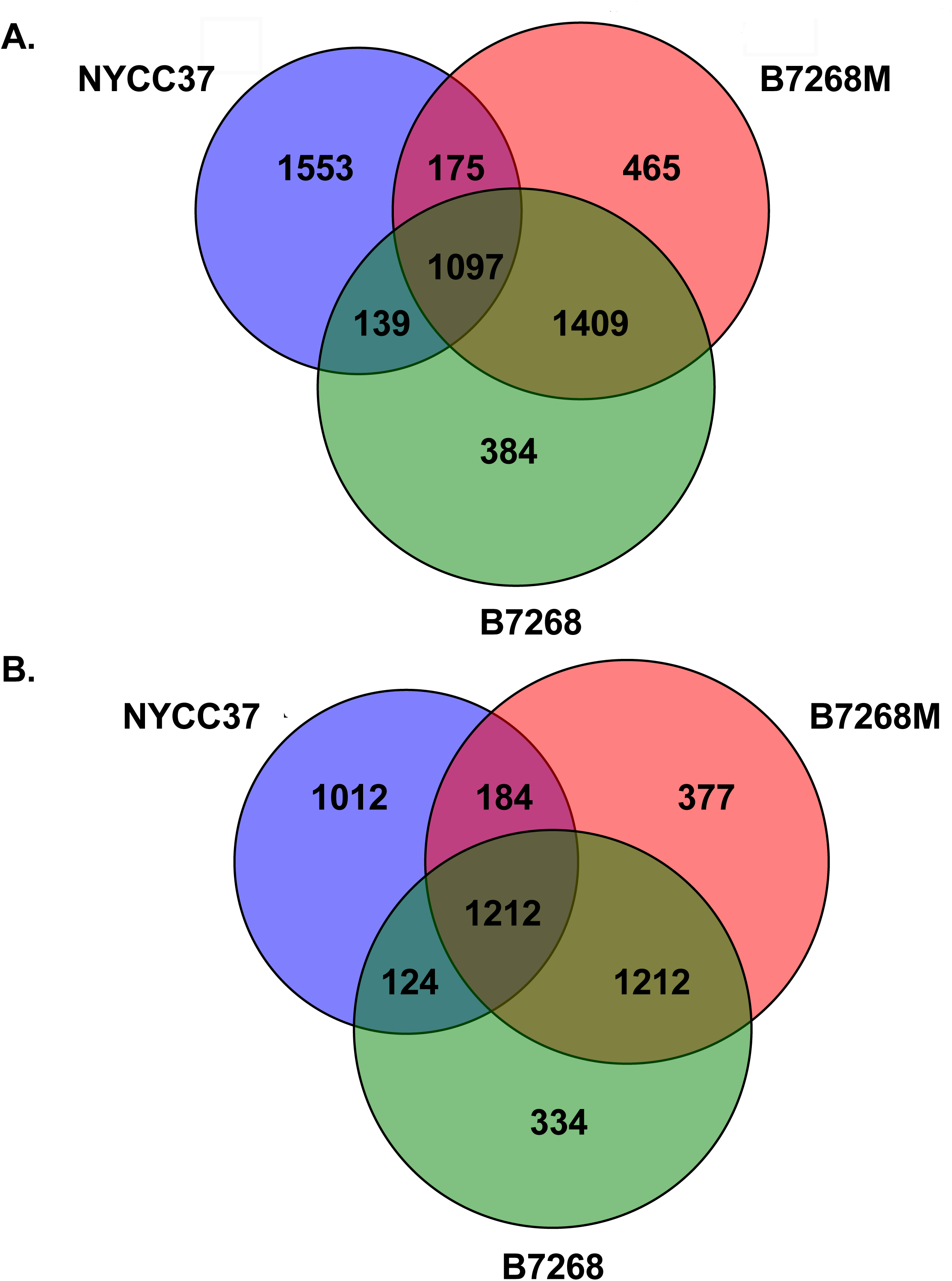
Differential gene expression in laboratory-derived and clinically resistant *T. vaginalis* versus sensitive lines. A. Venn diagram of number of genes in three Mz-resistant strains whose expression is up-regulated in comparison with nine Mz-sensitive strains. B. Venn diagram of number of genes in the same strains whose expression is down-regulated in comparison with nine Mz-sensitive strains. Changes in genes where difference in gene expression is significant to p<0.001 are shown. All the circles are sized based on marker numbers.

### Mz resistance is associated with regulation of genes involved in carbohydrate metabolism, hydrogenosome function, and oxygen detoxification

Considering the *T. vaginalis* gene expression data on their own, we found significant changes in expression of genes in carbohydrate metabolism, hydrogenosome function, and oxygen detoxification pathways. Major carbohydrate pathway genes, which have previously been implicated in Mz resistance [24, 25, 30, 39, 40] to be among the most differentially regulated in our RNA-Seq study (p < 0.05; **Additional file 6; Additional file 1, Figure S3**). Many genes coding for early stage glycolysis enzymes (*e.g.*, *phosphoglucomutas*e [TVAG_054830, TVAG_300510, TVAG_405900], *glucokinase* [TVAG_397250, TVAG_260790], *phosphofructokinase* [TVAG_462920], and *fructose-bisphosphate aldolase* [TVAG_038440, TVAG_360700]) were down-regulated in all three resistant isolates. In contrast, we observed up-regulation of expression of enzymes involved in later stages of the glycolytic pathway (examples include: *phosphoenolpyruvate carboxykinase* genes TVAG_139300 and TVAG_310250; *malate dehydrogenase* gene TVAG_495880, and *malic enzyme* genes TVAG_416100 and TVAG_412220). In particular, down-regulation of four *ADH* paralogs, including *ADH1* (TVAG_228780) [26] and up-regulation of six paralogs suggests a relationship of Mz resistance with ADH gene regulation generally (**Figure 6A**). Differential regulation of genes in amino acid metabolic pathways (*e.g.*, glycine, serine, and threonine metabolism) was also noted (**Additional file 1, Figure S3, Additional file 6**). Together, these changes suggest that important shifts in several metabolic pathways accompany Mz resistance.

**Figure 6.**
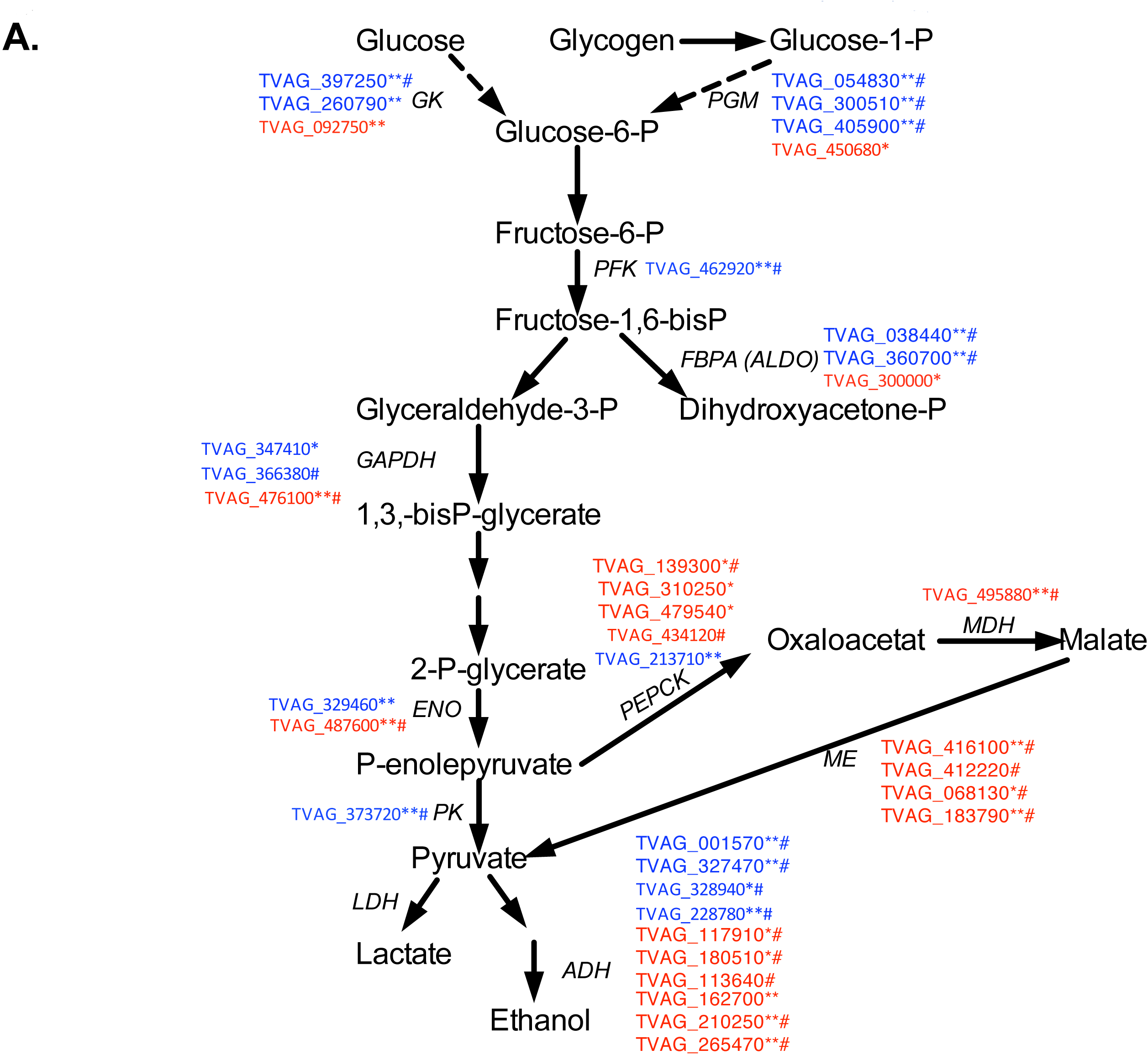
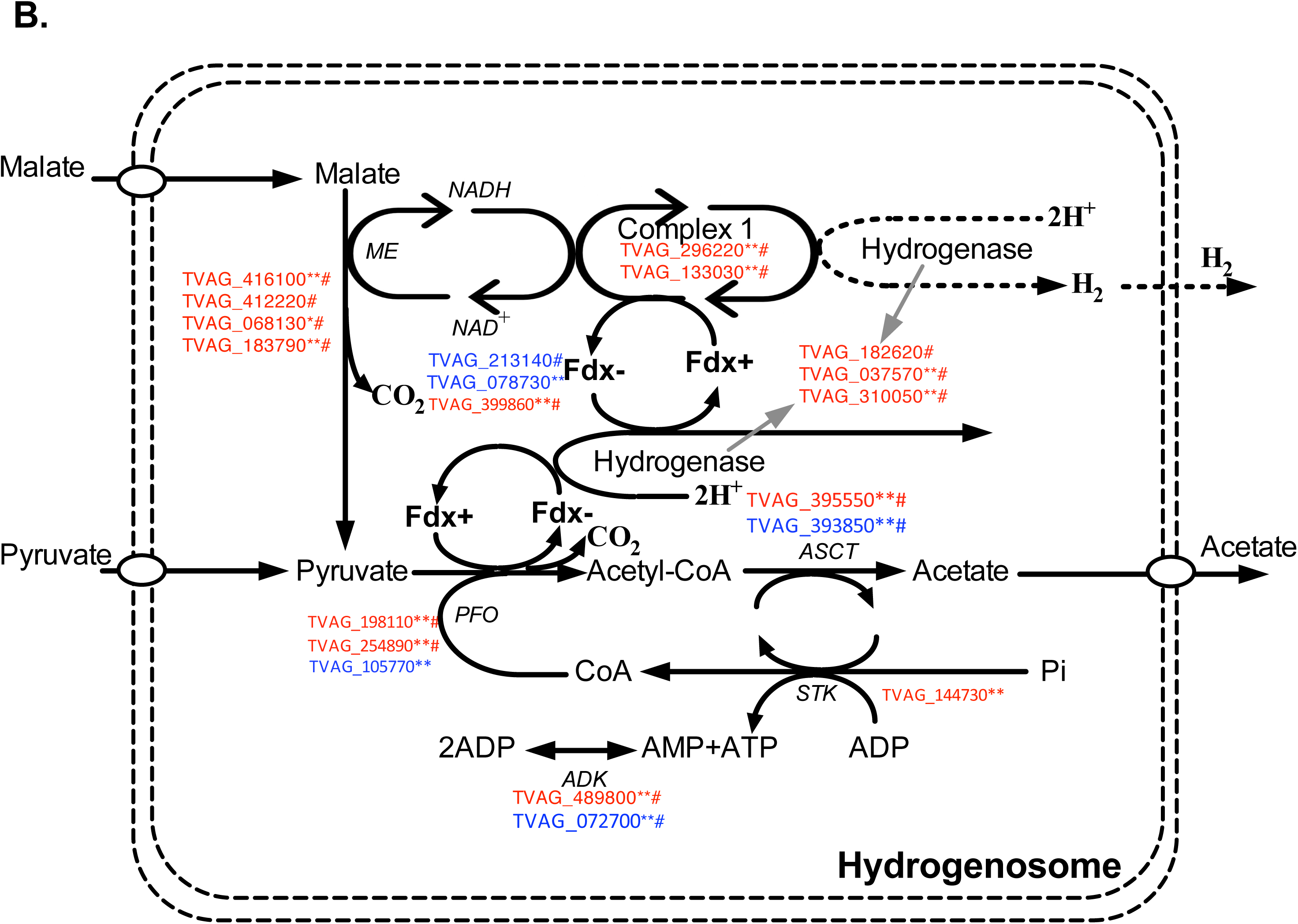
Differential gene expression of glycolytic and hydrogenosomal genes in laboratory-derived and clinically resistant *T. vaginalis*. A. Differential expression of genes in the (A) glycolytic pathway and (B) hydrogenosome (genes with products targeted to the hydrogenosome). Only genes with up or down regulated expression are shown as high-(red) and low-(blue) gene identifiers. * represents anaerobically resistant strains (B7268 and B7268-M), while # represents changes in clinically resistant strains. Enzyme abbreviations: *G*K, glucokinase; *PGM*, phosphoglucomutase; GPI, glucose phosphate isomerase; PFK, phosphofructokinase; *FBPA* (ALDO), fructose-1,6-bis-P aldolase; GAPDH, glyceraldehyde 3-P dehydrogenase; ENO, enolase; PEPCK, phosphoenolpyruvate carboxykinase; MDH, malate dehydrogenase; PK, pyruvate kinase; ME, malic enzyme; ADH, alcohol dehydrogenase; LDH, lactate dehydrogenase; *Fd*, ferredoxin; *PFO*, pyruvate:ferredoxin oxidoreductase; *ASCT*, acetyl:succinate coenzyme A transferase; *STK*, succinate thiokinase, *ADK,* adenylate kinase.

Of 179 predicted genes in the *T. vaginalis* G3 genome found as proteins in the hydrogenosome [41], 62 were down-regulated and 29 up-regulated in the whole transcriptome data of the three drug resistant isolates (B7268, B7268M, NYCC37) relative to the transcriptome data of our nine sensitive isolates. The majority of up-regulated genes have functions associated with metabolism and energy production in this organelle (**Figure 6B**), *e.g., PFO A* (TVAG_198110), an enzyme that reduces *ferredoxin (Fdx).* Down-regulation was primarily observed among hydrogenosome membrane proteins, such as iron-sulfur flavoproteins that play an important role as detoxifying reductases of Mz [42].

*T. vaginalis* oxygen tolerance pathways are likely important mediators of clinical resistance [22, 43, 44]. We observed changes in multiple genes associated with oxygen detoxification (**Additional file 61)**. For example, down-regulation in expression of paralogs of oxygen-regulating *thioredoxin* and *thioredoxin peroxidase* genes were observed in all three Mz resistant strains (**Additional file 6**), with the exception of *thioredoxin* paralogs TVAG_085330, and TVAG_428740 (which is in the top 3% of up-regulated genes ranked by expression level; **Additional file 6**). *Rubrerythrin* (TVAG_064490) showed increased expression in Mz resistant strains B7268 and B7268-M. Of the seven *flavin reductase* genes, *FR4* [TVAG_ 406950] was downregulated in all three Mz resistant strains, whereas *FR1* [TVAG_517010]) was absent in NYCC37 and B7268-M and is in the top 1% of down-regulated genes in the transcriptome data. We also observed transcriptional variation in paralogs of *superoxide dismutase* (*SOD;* TVAG_039980 down-regulated, TVAG_049140 up-regulated), a family of genes involved in scavenging oxygen reactive species as part of an oxygen defense mechanism in *T. vaginalis* previously shown to be involved in Mz resistance [24]. Finally, previous studies associated oxygen-insensitive *nitroreductase* (*Ntr*) and *nitroimidazole reductase* (*NimA*) gene mutations with Mz activation and inactivation, respectively [32, 45]. Our study supports the importance of those genes by detecting significantly lowered expression of four out of eleven copies of *ntr* genes [TVAG_455650 (*ntr 2*), TVAG_052580 (*ntr 5*), TVAG_354010 (*ntr 6*), TVAG_277870 (*ntr 10*)] in the transcriptome data of our three Mz resistant strains compared to the transcriptome data of the nine Mz sensitive strains. One paralog of *NimA* (TVAG_005890) was down-regulated in all three resistant strains, confirming previous studies [32]. A summary of the genes and their SNPs described in this section is shown in **Additional file 4**.

### Similar genetic changes associated with Mz resistance occur in two distantly related trichomonad species

To investigate whether there are evolutionarily common mechanisms of Mz resistance in trichomonads, we undertook whole genome shotgun sequencing and SNP detection of three strains of *Tritrichomonas foetus*, a trichomonad distantly related to *T. vaginalis* that also causes a venereal disease in cattle. We used Mz-sensitive KV1, and Mz-resistant lines KV1_M100 (exhibiting *in vitro* aerobic resistance) and KV1_1MR100 (exhibiting *in vitro* aerobic and anaerobic Mz resistance) derived by Kulda and colleagues from KV1 [46] (**Additional file 1, Figure S4**). Details of the sequencing are shown in **Table 1**. Sequenced reads of the KV1 genome were assembled into 194,695 contigs (N50=2,054 bp) with a size range of 61-19,994 bp. A BLASTX search of the KV1 assembly against the *T. vaginalis* G3 reference assembly returned 28,632 orthologs of *T. vaginalis* genes (data not shown). We identified 183 SNPs in the orthologs, 16 of which (in 16 orthologs) were present in both Mz-resistant *Tr. foetus* lines (**Figure 7** and **Additional file 7**). These 16 orthologs include genes encoding a sugar transporter (TVAG_015340), an actin-like protein (TVAG_371880), a BspA-like surface antigen (TVAG_228080), an integral membrane protein (TVAG_128430), a serine peptidase (TVAG_020790) notable for being downregulated in the transcriptome data of our three *T. vaginalis* Mz resistant strains compared to the transcriptome data of the nine *T. vaginalis* Mz sensitive strains, and a TPR domain protein (TVAG_371930) that was upregulated in two. The remaining 10 orthologs all encoded conserved hypothetical proteins. Another potentially interesting set of SNP-containing orthologs comprised those that were observed in a resistant *Tr. foetus* strain and were also up- or down-regulated in all three resistant *T. vaginalis* lines in our RNA-Seq analysis. We found three such orthologs in KV1_M100 and 17 in KV1-MR100. One of them, an upregulated gene encoding *pyruvate:ferredoxin oxidoreductase A* (*PFO A*, TVAG_198110), also shared a nonsynonymous SNP in two of three lab-derived resistant *T. vaginalis* lines.

**Figure 7.**
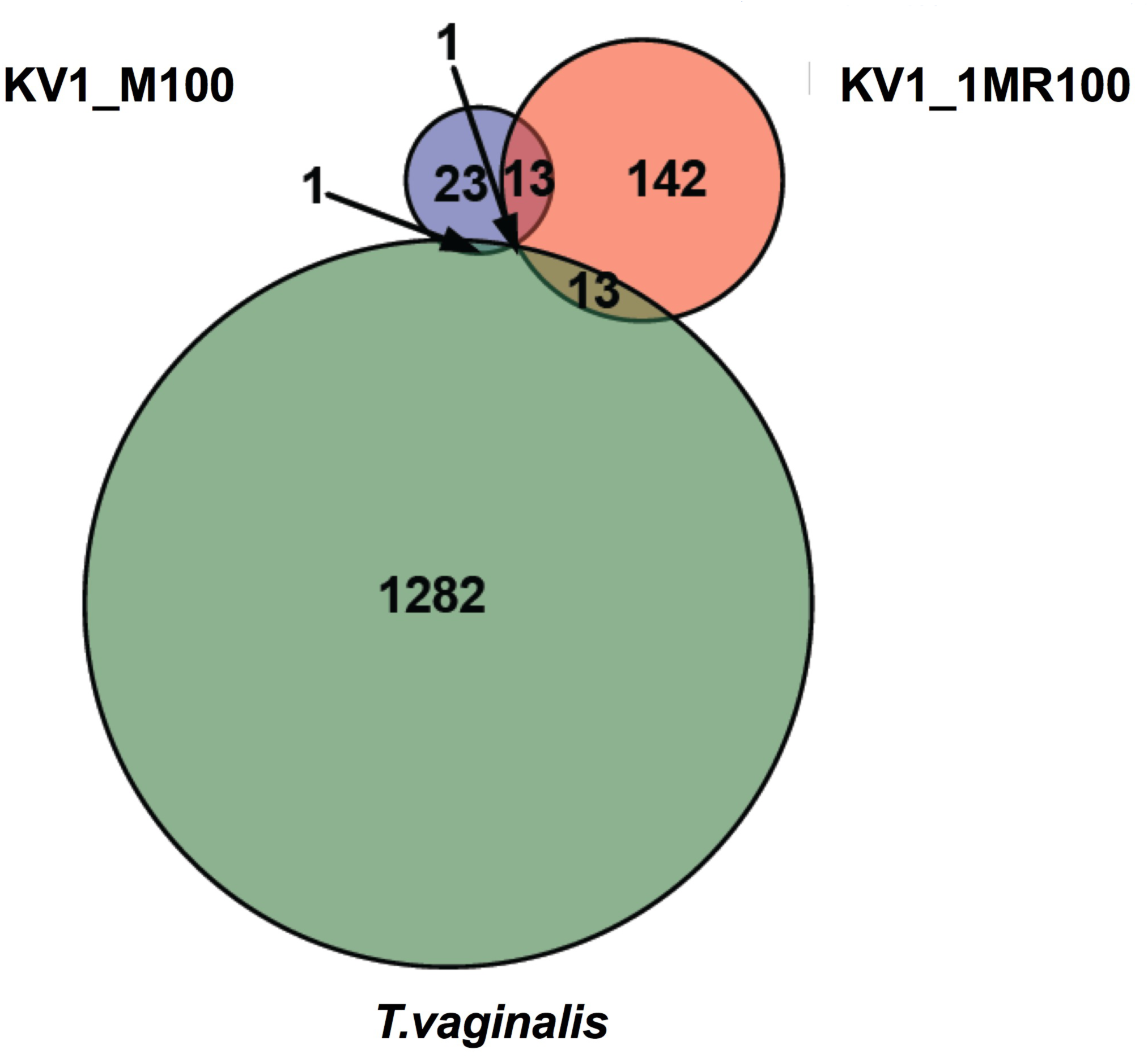
Common genetic changes in laboratory-derived *T. vaginalis* and *Tr. foetus* lines. Venn diagram of number of genes with nonsynonymous mutations in two laboratory-induced Mz resistant strains of *Tr. foetus*. Each resistant strain (KV1_M100- *in vitro* aerobic resistance and KV1_1MR100- *in vitro* aerobic and anaerobic Mz resistance) was compared to its ancestral sensitive strain (KV1). Comparison is also made with the three *T. vaginalis* derived lines B7008M, B7268M, F1623M. Circles are in proportion to the numbers of SNPs found in the genes.

**Table 1.** Characteristics of whole genome shotgun sequencing datasets of three *Tr. foetus* strains, KV1, KV1_M100, and KV1_1MR100. Coverage was determined using the formula N x L/G, where N = number of reads, L = average length of reads, and G = genome size.

|  | *Tr. foetus*<br>KV1 | *Tr. foetus*<br>KV1_M100 | *Tr. foetus*<br>KV1_1MR100 |
| --- | --- | --- | --- |
| **Library insert size** | 350 bp | 350 bp | 350 bp |
| **Read length** | 100 nc | 100 nc | 100 nc |
| **Total no. reads** | 67,088,722 | 75,792,262 | 70,337,248 |
| **Coverage** | 44.27 x | 46.23 x | 49.48 x |
| **Avg. GC%** | 30 | 30 | 30 |
| **Assembly** | Velvet | Not done | Not done |
| **No. contigs** | 194,695 | - | - |
| **Contig N50** | 2,054 bp | - | - |
| **No. orthologs with**<br>***T. vaginalis*** | 28,362 | - | - |

## Discussion

Our study of genetic variation and gene expression related to drug resistance in multiple *T. vaginalis* isolates is the most comprehensive to date and offers a new baseline for functional and biochemical studies of this organism. We used ddRAD-Seq to genotype unique regions of the highly repetitive *T. vaginalis* genome, revealing it to be a useful method that can be applied to other protists whose genomes are refractory to cost-effective whole genome sequencing. Using the dataset of 3,923 SNP markers, we explored the genetic diversity of *T. vaginalis* and present a picture of the parasite’s population structure and linkage disequilibrium (LD) at higher resolution than previous studies [14, 36]. This dataset represents a high confidence set of genetic variants, and greatly expands upon the limited number of genetic markers available for this parasite. Our study also offers a panel of biomarkers that could be used to advance personalized treatment and prevention of trichomoniasis.

Genetic variation and exchange through recombination is important to consider in parasites as it influences the spread of drug resistance and virulence genes among them. Our study confirms the existence of two genetically distinct *T. vaginalis* populations [13, 14, 36, 45], congruent with sexually reproducing organisms [34]. Genome-wide LD, reported from analysis of a small number of genes in previous studies [13, 14], was found to decay rapidly within 1-5 kb in both populations, providing strong evidence for recent genetic exchange. Differences in LD decay between the two populations suggests that different degrees of inbreeding and rates of transmission might be responsible for maintenance of the two-type population structure. Based on previous work [14, 45], we also expected to observe a difference in average drug susceptibility between the two *T. vaginalis* populations. Higher mean values of Mz resistance were found in one population compared to the other (98.5 µg/ml *vs.* 126.2 µg/ml), although the difference was not statistically significant.

A primary goal of generating the SNP markers in this study was to undertake a genome-wide association analysis to identify novel genetic indicators of Mz resistance. From analysis of ~100 global *T. vaginalis* Mz resistant and sensitive strains, we identified 72 SNPs associated with moderate or high Mz resistance, which represent a panel of biomarkers that could be used to advance personalized treatment and prevention of trichomoniasis. Currently only a single “reflex test” that identifies a mutation in the *T. vaginalis ntr6* gene by real-time PCR (http://www.prnewswire.com/news-releases/medical-diagnostic-laboratories-llc-announces-a-complimentary-reflex-test-to-determine-metronidazole-resistance-in-trichomonas-vaginalis-145736975.html) is available for clinical testing of resistance. Of the 72 biomarkers, several (*e.g.,* those found in TVAG_124910 [conserved hypothetical] and TVAG_197160 [CAMK family protein kinase]) were also recurrently identified in genes in other analyses, such as a comparison of three laboratory-derived Mz resistant lines with their isogenic parental strains. While we were unable to undertake a direct functional analysis of these new putative resistance mutations and genes because of the limitations of *T. vaginalis* transfection, whole transcriptome analysis of three resistant and nine sensitive isolates showed several of the mutations to be associated with the functional consequence of a change in gene expression.

A major finding of this study is changes in expression of metabolic pathway genes in Mz resistant parasites. For example, genes coding for early stage glycolysis enzymes were found to be down-regulated in drug resistant isolates relative to sensitive isolates, compared to up-regulation of expression of enzymes involved in later stages. Gene expression of proteins located in the hydrogenosome identified 62 as down-regulated (mainly involved in hydrogenosome metabolism), and 29 as up-regulated (mainly hydrogenosomal membrane proteins), in drug resistant isolates relative to sensitive isolates, including *PFO A* (TVAG_198110), an enzyme that reduces *ferredoxin (Fdx).* Down-regulation of genes in the *PFO-ferredoxin* pathway in hydrogenosomes has been hypothesized to induce anaerobic drug resistance [44, 47, 48]. Our study revealed down-regulation of one *PFO* gene (TVAG_105770) specific to anaerobic resistance, as well as both up-regulation and down-regulation of *Fdx* genes, thus suggesting their potential involvement in Mz resistance, but not a crucial or anaerobic resistance-specific role. Anaerobic resistance has also been linked to the thioredoxin reductase (*TrxR)* pathway, which is essential for cell growth and protection of cells during oxidative stress [24]. Under low- or no-oxygen conditions, *TrxR* in the cytosol reduces and creates covalent adducts with Mz, effecting multiple reactions in the cell. *TrxR* in our study was expressed similarly in all strains and thus seems to be regulated only at the protein level, which has been shown in a previous study [24]. However, expression changes in *Trx, superoxide dismutase* (*SOD*) and *tryparedoxin peroxidase (TxnPx)* support an important role for thiols in response to Mz treatment, which maybe as a result of Mz resistance rather than a cause as suggested by Leitsch *et al.,* [24], and furthermore seem to be important in both aerobic and anaerobic resistance. Our results, however, present opposite expression patterns from studies for *SOD*, *TxnPx,* and *Trx* genes [24], possibly due to experiments being performed in different strains, or under anaerobic conditions, and analysis of protein rather than RNA levels. Finally, although our research suggests that similarly to *T. vaginalis*, anaerobic resistance in *Tr. foetus* shares mutations in enzymes related to hydrogenosomal processes, our knowledge of genetic mechanisms involved in aerobic vs. anaerobic adaptations is limited by insufficient orthologous gene information for *Tr. foetus*, and more work regarding those mechanisms will be necessary.

An important question arising from this work is what our analyses reveal about the evolution of Mz resistance in *T. vaginalis* or other trichomonads. We observed mutations in genes shared between several genetically distinct laboratory-derived *T. vaginalis* lines and >100 *T. vaginalis* clinical isolates, as well as with the distantly-related *Tr. foetus* that causes venereal disease in cattle, suggesting strong natural selection on the same sets of genes and the presence of shared pathways of resistance. We propose that one of the first means for developing Mz resistance in trichomonads is the development of tolerance to high oxygen levels, which provides a protective mechanism against the drug. For example, our transcriptome data support a key role of *ferredoxin reductase 1* (*FR1*), and extend this role to other genes involved in oxygen-scavenging. The importance of *FR1* is in agreement with Leitsch *et al.,* who first showed reduced FR1 activity in six resistant *T. vaginalis* strains [24, 26], and restoration of Mz sensitivity in resistant B7268 by transfection and expression of *FR1* [23]. *T. vaginalis* already harbors mechanisms to cope with a highly fluctuating oxygen environment [44], which may make it “pre-adapted” to Mz treatment. This hypothesis is further supported by the extremely rapid development of aerobic resistance under lab conditions (<50 *in vitro* passages), and the emergence of clinical Mz resistance within only two years of introduction of the drug [49]. Other genes may play an important role in developing subsequently higher levels of resistance. For example, several genes in highly resistant isolates show the same mutation recurrently appearing; examples include disulfide oxidoreductase (TVAG_049830), chromodomain helicase DNA binding protein (TVAG_302840), 5'->3' exoribonuclease (TVAG_424140), and numerous other hypothetical proteins whose function has yet to be determined (**Additional file 4**). This is in contrast to lab-induced anaerobic resistance which in both *T. vaginalis* [19] and *Tr. foetus* [46] appear to show stepwise selection over a period of time, most likely proceeding through recurrent mutations in a limited number of genes, at which point resistance "stalls" while new mutations emerge. A similar phenomenon was previously reported in stepwise antibiotic resistance in bacteria [50, 51]. While the details of these mechanisms and their fitness costs should be further explored, we provide here a first insight into adaptation of the unicellular eukaryote *T. vaginalis*.

## Conclusion

We have identified a panel of genetic markers associated with *T. vaginalis* resistance to metronidazole. Maintaining redox homeostasis seems to be a major problem that parasites face while adapting to Mz treatment and its disruption can shift multiple biochemical processes in the cell. These multiple pathways described here work together in facilitating Mz resistance. Changes in genes and pathways relevant for drug resistance that were shared between clinical isolates and lab-derived strains suggest conservation of the genetic pathways through which resistance occurs. Finally, in *Tr. foetus*, a distantly related species that causes an analogous disease in cattle, we observed similar changes in several derived drug resistant lines.

## Materials and Methods

### Parasite strains, culturing and DNA extraction

We used a set of 102 *T. vaginalis* isolates originating from eight countries: Australia (13), Italy (2), Mexico (8), Papua New Guinea (18), South Africa (9), Mozambique (1), United States (50), and the United Kingdom (1) (**Additional file 2**). Most of the strains were isolated from women undergoing routine examination in sexual health clinics and then adapted to growth in culture. Additional isolates included the reference lab strain G3, and three *in vitro*-derived Mz resistant lines and their isogenic parents, whose Mz resistance was selected for using the method of Kulda *et al*., [21] (see **Figure S2** in **Additional file 1)**: Mz-sensitive parental strains F1623 and B7708 and their Mz-resistant derivatives F1623-M and B7708-M (MLC > 400 ug/ml, anaerobic) [19], and moderately resistant parental strain B7268 (MLC > 200 μg/ml) [29] and its derivative B7268-M (MLC > 400 ug/ml, aerobic and anaerobic) [38]. All three derived lines are stably resistant in the absence of drug pressure. We also obtained three *Tritrichomonas foetus* lines derived using a similar selection method [47]: *Tr. foetus* KV1 is the Mz-sensitive parental strain to two resistant lines: KV1_M100 (exhibits both *in vivo* and *in vitro* aerobic resistance) and KV1_1MR100 (exhibits *in vitro* anaerobic resistance only), respectively (see **Figure S4** in **Additional file 1**). *T. vaginalis* and *Tr. foetus* isolates were cultured in modified Diamond's media as described [12], pelleted by centrifugation, and parasite DNA extracted using a standard phenol-chloroform procedure [14].

### Metronidazole susceptibility phenotyping

*T. vaginalis* susceptibility to Mz was determined using a standard minimum lethal concentration (MLC) protocol under aerobic conditions [52]. Each isolate was grown in culture media in 96 well plates in the presence of serial dilutions of Mz ranging from 0.2 to 400 μg/ml for 48 hours. The lowest concentration at which no motile parasites were observed by microscopy was recorded as the MLC. Each assay was repeated two times. Control strains (CDC 252, resistant to >400 μg/ml Mz, and Mz sensitive CDC 520 [53]) were obtained from the Centers for Disease Control and Prevention (CDC) and used in each assay. Low-level resistance was defined as aerobic MLC 50–100 µg/ml, moderate-level resistance as 100-400 µg/ml, and high-level resistance as 400 µg/ml or greater, as per previous studies [17, 54].

### *In silico* restriction enzyme optimization for ddRAD sequencing

We used ddRAD-Seq to sample unique regions of the highly repetitive *T. vaginalis* genome. We first evaluated *in silico* digests of the reference assembly of strain G3 using several common choices of restriction enzymes implemented in GMBX_digest_v1.0.pl script (https://github.com/GenomicsCoreLeuven/GBSX) and the SimRAD package in R [2], starting with digestion with six-base cutting site enzymes (EcoRI, PstI, SdaI, SgrAI, SbfI) to determine the best enzyme for cutting within unique regions of the genome (defined as regions free of highly repetitive TEs or genes in multicopy families, based on the current genome annotation (www.Trichdb.org). EcoRI was found to cut most frequently in the unique regions of the genome (**Figure S5A** in **Additional file 1**). Subsequent testing of EcoRI in combination with other restriction enzymes, both experimentally (data not shown) and *in silico*, revealed that a double digest of EcoRI and NlaIII produced the optimal 650-850 bp fragment size distribution for Illumina library preparation and sequencing (**Figure S5B** and **Figure S5C** in **Additional file 1**).

### ddRAD library preparation and Illumina sequencing

ddRAD sequencing libraries were prepared using EcoRI and NlaIII digested DNA followed by the protocol described in [55]. Sequencing adapters containing a combinatorial in-line barcode and standard Illumina multiplexing read index were used to individually barcode samples such that the identity of each isolate was kept intact. Libraries were sequenced on an Illumina HiSeq 2000 following standard methods for cluster generation and sequencing, and paired end (PE) reads of 100 bp were generated. Samples were de-multiplexed and individual FASTQ files obtained using Google Documents spreadsheets and database tools in Python [55].

### SNP discovery and filtering

To estimate the SNP false discovery rate and optimize SNP filtering, the Perl script IRMS.pl [56] was used to introduce 99 random SNPs into the reference G3 genome *in silico*. SNPs were then detected by aligning ddRAD sequencing G3 data to the G3 reference with introduced SNPs using BWA version 5.09 to align reads [57] and GATK to call variants [58]. Multiple filtering steps included removing SNPs based on quality (QUAL>30), approximate read depth (DP<5), minimal genotype quality (GQ<25), quality by depth (QD< 5) and minor allele frequency (MAF) <0.05. We recovered 82% of introduced SNPs upon alignment with the ddRAD re-sequenced G3 data, thus the same criteria were applied to our actual ddRAD dataset (**Figure S6** in **Additional file 1**).

Illumina PE reads generated for 102 isolates were aligned to the G3 reference genome in the TrichDB database [59] using BWA. BAM alignment files of all sequenced isolates were merged and locally realigned to the reference assembly using GATK RealignerTargetCreator in order to minimize the number of mismatching bases across all the reads [58]. Genotypes were called from merged BAM files using GATK. In addition to the above-mentioned filtering criteria, SNPs detected from multiple samples were further filtered by removing those with >40% missingness, and those found within TEs. These filtering steps produced a final set of 3,923 high quality filtered SNPs from the 191,671 total SNPs identified from 102 *T. vaginalis* libraries. Direct SNP comparisons between parental and drug resistant derived pairs were made for each SNPs such that those SNPs found in ancestral pair must match with reference G3 strain (sensitive), and must differ from derived stains with at least 3 reads for each SNP.

### Sanger sequencing for SNP validation

Using the reference G3 assembly in the TrichDB database [59], primers were designed to 150-300bp regions flanking six of the 72 SNPs associated with MLC > 100 μg/ml and used to amplify DNA from 20-24 *T. vaginalis* isolates (**Table S1 in Additional file 1**). PCRs were performed in 25 µL reactions containing ~20 ng genomic DNA, 0.2 µM of each primer, 200 µM of each dNTP, 1.5 mM MgCl_2_, 1 U of Taq DNA polymerase (Sigma) and 1× PCR buffer. Standard PCR amplification programs were used with an initial denaturation for 5min and 30 cycles, and PCR products visualized on 1.5% agarose gels. Amplicons were cleaned with QIAquick PCR purification kit (Qiagen) and sequenced using standard Sanger sequencing chemistry. The identity of ~73% (range ~63% to 88%) of the SNPs identified through ddRAD sequencing were confirmed by Sanger sequencing.

### Population structure

Population structure was assessed from the 3,923 high-quality SNPs representing 100 isolates using Discriminant Analysis of Principal Component (DAPC) implemented in the R *adegenet* package, using “average” in the *snpzip* function [37, 60]. *snpzip* uses DAPC to calculate the contribution (loadings) of each SNP to each population group or phenotype. Correction of population structure was performed by regressing along the first PCA axis. DAPC with the *snpzip* function and centroid clustering was also used to identify SNPs that distinguish Mz-resistant and Mz-sensitive isolates. SNP effects from the final data set were predicted using SnpEff [61], and functional categories were determined using the Kyoto Encyclopedia of Genes and Genomes (KEGG) [62, 63] metabolic enrichment tools in TrichDB [59].

### Linkage disequilibrium

After filtering out rare variants (MAF <0.05) which could give spurious LD, we recovered 3,894 SNPs from the 3,923 SNPs used to determine population structure. Filtering this set within each population gave 2,837 SNPs covering 898 genomic contigs in one population, and 2,694 SNPs covering 872 contigs in the other. Linkage disequilibrium decay from SNPs was determined using the function r^2^ from the toolset PLINK (http://pngu.mgh.harvard.edu/purcell/plink), with pairwise r^2^ only considered within each contig. The decay of LD over distance was estimated using the Hill and Weir formula [64], and a non-linear model was used in order to fit the data to the decay function.

### RNA-seq of *T. vaginalis* Mz resistant and sensitive parasites

Three Mz-resistant *T. vaginalis* isolates B7268, B7268-M, and NYCC37, and nine Mz-sensitive isolates (G3, GOR69, NYCA04, NYCB20, NYCD15, NYCE32, NYCF20, NYCG31, SD2) were grown overnight in 14 ml sealed tubes and total RNA was isolated using the Qiagen® RNeasy Mini Kit, including a column DNase treatment using the Qiagen® RNase-Free DNase kit. Polyadenylated RNA was purified from 5 μg of total RNA using the Dynabeads® mRNA DIRECT^TM^ Purification Kit. All experiments were carried out in triplicate. First strand synthesis was carried out by mixing the entire fraction of isolated polyA+ RNA (8 μl) with 0.5 μl Random Primers (3 ug/μl, Invitrogen), 10 mM DTT and 0.25 μl ANTI-RNase (15-30 U/μl, Ambion) in 1x First-Strand Synthesis Buffer (5x, Invitrogen), with incubation at 65°C for 3 minutes to remove RNA secondary structures, then placed on ice. A total of 0.5 μl SuperScript® III enzyme (200 U/μl, Invitrogen) and dNTPs to a final concentration of 0.125 mM were added to the mixture and reverse transcription was carried out using the following incubations: 25°C for 10 minutes, 42°C for 50 minutes, 70°C for 15 minutes. The resulting cDNA/RNA hybrid was purified from the mix using Agencourt RNAClean XP Beads according to manufacturer's instructions. Second-strand synthesis was carried out by mixing the purified cDNA/RNA hybrid with 1 μl dUTP mix (10 mM, Roche), 0.5 μl Ribonuclease H (2 U/μl, Invitrogen). 1 μl DNA polymerase I (5-10 U/μl, Invitrogen) in 1 X NEB buffer 2 with 2.5 mM DTT. This mixture was incubated at 16°C for 2.5 hours. The resulting double stranded cDNA was purified using Agencourt AMPure XP beads according to manufactures instructions. The cDNA was end repaired by mixing 5 μl T4 DNA Polymerase (3 U/ μl, NEB), 2 μl Klenow DNA Polymerase (3-9 U/μl, Invitrogen), 5 μl T4 Polynucleotide Kinase (10 U/μl, NEB) in 1 x T4 DNA Ligase buffer with 10 mM ATP (NEB) with 0.4 mM dNTPs. The mixture was incubated at room temperature for 30 minutes. The end-repaired cDNA was purified using Agencourt AMPure XP beads according to manufactures instructions. The purified end-repaired cDNA was then taken through A-tailing, adapter ligation and PCR enrichment using the Illumina TruSeq Stranded mRNA Sample Preparation Kit with different barcodes for each sample. The libraries were pooled and sequenced on the Illumina HiSeq 2500 with 101 cycles, paired-end reads, and multiplexing.

### Transcriptome analyses

Approximately 10-20 million reads were generated per RNA-seq library and all RNA-seq data sets were aligned to the G3 reference genome sequence [59] using BWA version 5.09 [57] with 1300 bp used as a distance between the reads, yielding an overall alignment rate > 95% for all libraries. Counts of reads for each gene were obtained using the program HTseq [65]. Expression patterns in the three Mz resistant (B7268, B7268-M and NYCC37) strains and the nine (G3, GOR69, NYCA04, NYCB20, NYCD15, NYCE32, NYCF20, NYCG31, SD2) sensitive strains were compared pair-wise against the reference sensitive strain G3 using the DESeq2 R package [66]. Differences in gene expression between G3 and the sensitive strains were subtracted from the resistant strain differences. Quality control of the data was undertaken by comparing all three replicates of each sample. The raw read counts were normalized by *rowMeans* normalization function based on counts. A statistical test (*binomialWaldTest*) was used to detect up- and down-log_2_ fold changes in gene expression. Differential gene expression was explored using the *MAplot* function in *DESeq2* R package, p-values were adjusted using false discovery rate (FDR), and only changes where padj <0.01 were considered significant.

### Metabolic pathway analysis

Pathway enrichment analyses were performed using the TrichDB KEGG metabolic pathway tool using a P-value cutoff of 0.05 [59].

### *Tr. foetus* whole genome sequencing

Whole genome sequences of three *Tr. foetus* samples (1) KV1 (Mz sensitive), (2) KV1_M100 (derived *in vitro* from KV1 and exhibiting aerobic Mz resistance), and (3) KV1_1MR100 (derived *in vitro* from KV1 and exhibiting aerobic and anaerobic Mz resistance) as described in Ref. [46], were generated by standard Illumina sequencing of 350 bp and 500 bp insert libraries and generation of 100 bp paired-end reads. *Tr. foetus* KV1 Illumina sequencing reads were assembled *de novo* into 194,695 contigs (N50=2,054 bp) with a size range of 61-19,994 bp using Velvet [67]. Sequencing produced between 44.3x coverage and 49.5x coverage of each isolate [67]. Illumina reads from KV1, KV1_M100 and KV1_1MR100 were aligned against the KV1 contigs and SNPs were called using the same GATK pipeline as described above for *T. vaginalis* SNP detection. We identified 627 high quality SNPs, of which 418 were found in KV1_1MR100, 86 were found in KV1_M100, with 97 were shared between these two resistant lines. To annotate SNPs we used BLASTX [68] (with -max_target_seqs 1 - culling_limit 1 -seg no) to search a local *T. vaginalis* protein database and identify orthologs genes. Only BLASTX hits >50 bp length were considered valid and received a *T. vaginalis* gene attribute. Candidate SNPs were then assigned to the appropriate *T. vaginalis* orthologs.

## Data access

All sequence data have been deposited in the Sequence Read Archive of GenBank with the accession numbers SRP057357 (*T. vaginalis* ddRAD data, whole genome sequencing data and RNA-Seq data) and SRP057311 (*Tr. foetus* whole genome sequencing data).

## Abbreviations

DAPC: discriminant analysis of principal component; LD: linkage disequilibrium; Mz: metronidazole; PCA; principal component analysis; SNP: single nucleotide polymorphism; TE: transposable element.

## Competing interests

The authors declare that they have no competing interests.

## Authors' contributions

MB designed the study, performed experiments, analyzed the data, and drafted the paper. SW made the RNA-seq libraries. GT performed Sanger sequencing validation experiments. PS helped with ddRAD sequencing experimental design. WS provided lab lines and training in drug resistance assays. PH, TC, CL and PT performed NGS sequencing and raw data processing of three whole genomes. SS undertook data checking, proofreading, and writing the manuscript. JC oversaw all aspects of the research and experimental design, as well as manuscript writing.

## Acknowledgments

We thank members of the Carlton lab for critical reading of the manuscript and Jonathan Flowers and Swapna Uplekar for helpful discussions. We also thank Peter Augostini at CDC for providing training and protocols for the minimum lethal concentration phenotypic assay. MB is partially supported by R01 AI097080 to P. Kissinger (Tulane University), S.D.W. by the MacCracken Program in the New York University Graduate School of Arts and Science and a Fleur Strand Fellowship, G.E.T. by a New York University Gallatin Undergraduate Research Fund award, and PT by the Chang Gung Memorial Hospital Research Fund (CMRPD3D0181-3). The findings and conclusions in this report are those of the authors and do not necessarily represent the official position of the Centers for Disease Control and Prevention. None of the authors have any compete

## Additional Files

**Additional file 1. Supplementary figures and tables:** Table S1. List of SNPs validated by Sanger sequencing. **Figure S1.** Summary of analysis undertaken in this study. **Figure S2.** Schematic representation of *in vitro*-induced Mz resistance in three *T. vaginalis* isolates, as described in [19, 29, 38]. **Figure S3.** KEGG metabolic pathways coded by genes up or down regulated in transcriptome data from three drug resistant *T. vaginalis* isolates (B7268, B7268M, NYCC37) relative to nine sensitive strains. A. Common down-regulated metabolic pathways. B. Common up-regulated metabolic pathways. Only metabolic pathways with Bonferroni corrected significance of P < 0.05 are shown. **Figure S4.** Schematic representation of how the two *in vitro*-derived Mz resistant isolates KV1-M100 and KV1-1MR100 of *T. foetus* were generated from KV1, as described in reference [46]. **Figure S5.** *In silico* determination of ddRAD restriction enzyme sites, fragment sizes, and number of fragments. A. Number of restriction sites in the G3 reference strain for five restriction enzymes. The x-axis represents the restriction enzyme, and the y-axis the log_10_ of number of the sites calculated as existing in either repetitive or unique regions of the genome. **B.** Fragment size and number for EcoRI and NlaIII ddRAD enzyme pairs. **C.** Fragment size distribution based on the G3 sequence. **Figure S6.** Flowchart for read alignment and SNP discovery.

**Additional file 2. List of 102 *T. vaginalis* isolates used in the study.** The list includes the name of each isolate, its specific geographical origin, the world region to which it is assigned for population studies, the provider of each strain, the year it was isolated from a patient and the population type to which it clustered, the measured metronidazole MLC of the isolate, and references to publications in which the isolate has been included. NK: not known. *Antenatal samples from U.S. Navy wives or active duty U.S. Navy women.

**Additional file 3. SNPs associated clinically resistant *T. vaginalis* isolates.** SNPs associated with moderate Mz resistance (MLC ≥ 100) and high (MLC > 400) Mz resistance phenotype are shown. Yellow marks SNPs associated with both moderate and high-level resistance.

**Additional file 4. Summary of results for candidate genes with potential role in Mz resistance. Additional file 5. Comparison between drug sensitive *T. vaginalis* isolate and its resistant pair.** A. SNPs common to all three founder/derivative pairs (B7268/B7268-M, F1623/F1623-M, and B7708/B7708-M). B.SNPs common to two founder/derivative pairs: B7268/B7268-M and F1623/F1623-M. C.SNPs common to two founder/derivative pairs: B7708/B7708-M and F1623/F1623-M.D. SNPs common to two founder/derivative pairs: B7268/B7268-M and B7708/B7708-M. E. SNPs observed in a single founder/derivative pair: B7268/B7268-M. F. SNPs observed in a single founder/derivative pair: B7708/B7708-M. G. SNPs observed in a single founder/derivative pair: F1623/F1623-M.

**Additional file 6. Gene expression summary of comparison between B7268, B7268M, NYCC37 and nine sensitive strains. Table S3.** Down-regulated genes in B7268 resistant strain in comparison to nine sensitive strains.**Table S4**. Down-regulated genes in B7268M resistant strain in comparison to nine sensitive strains. **Table S5.** Down-regulated genes in NYCC37 resistant strain in comparison to nine sensitive strains. **Table S6.** Up-regulated genes in B7268 resistant strain in comparison to nine sensitive strains. **Table S7.** Up-regulated genes in B7268M resistant strain in comparison to nine sensitive strains. **Table S8.** Up-regulated genes in NYCC37 resistant strain in comparison to nine sensitive strains. **Table S9.** Commonly down-regulated genes in all resistant samples (B7268, B7268M, NYCC37) in comparison to nine sensitive strains. **Table S10**. Commonly up-regulated genes in all resistant samples (B7268, B7268M, NYCC37) in comparison to nine sensitive strains. **Table S11**. List of commonly down-regulated and up-regulated enriched metabolic pathways in all the samples. **Table S12**. Down-regulated genes present only in B7268 and B7268M. **Table S13**. Down-regulated genes present only in B7268 and NYCC37. **Table S14**. Down-regulated genes present only in B7268M and NYCC37. **Table S15**. Up- regulated genes present only in B7268 and B7268M. **Table S16**. Up-regulated genes present only in B7268 and NYCC37. **Table S17**. Up-regulated genes present only in B7268M and NYCC37. **Table S18.** List of down-regulated and up-regulated enriched metabolic pathways in B7268 and B7268M. **Table S19.** List of down-regulated and up-regulated enriched metabolic pathways only in B7268M.

**Additional file 7. Comparison between drug sensitive *T. vaginalis* isolates and their resistant pairs in *Tr. foetus*. A.** Changes common to both Mz resistant lines KV1_M100 and KV1_1MR100 when compared with Mz sensitive KV1 are shown, as well as changes unique to KV1_M100 and KV1_1MR100. Rows indicated in color represent shared SNPs with *T.vaginalis* lab derived resistance strains**. B.** List of enriched metabolic pathways containing SNPs identified in *T. foetus* KV1_M100 and KV1_MR100 lines.

